# Comparative transcriptomic and lipidomic analysis of fatty acid accumulation in three *Camellia oleifera* varieties during maturation

**DOI:** 10.1101/2024.03.07.583840

**Authors:** Dayu Yang, Rui Wang, Hanggui Lai, Yongzhong Chen, Yimin He, Chengfeng Xun, Ying Zhang, Zhilong He

**Affiliations:** Research Institute of Oil Tea Camellia, Hunan Academy of Forestry, Changsha 410004, China; National Engineering Research Center for Oil-Tea Camellia, State Key Laboratory of Utilization of Woody Oil Resource, Hunan Academy of Forestry, Changsha 410116, China; School of Tropical Agriculture and Forestry, Hainan University, Haikou 570228, China

**Author notes:** These authors contributed equally to this work (Dayu Yang, Rui Wang, Hanggui Lai).

**Keywords:** *Camellia oleifera*, Fatty acids, Lipid biosynthesis, Feedback regulation, Transcriptome analysis

## Abstract

*Camellia oleifera* stands out as one of China’s leading woody oil crops, famed for its tea oil, which boasts a high content of unsaturated fatty acids. Often referred to as "liquid gold" or "Oriental olive oil," this oil is prized for its nutritional benefits and culinary versatility, marking Camellia oleifera as a valuable agricultural and economic resource. This study provides an integrated investigation of the transcriptome and lipidome within seeds at the maturation stage across three *C. oleifera* varieties, revealing a significant relationship between fatty acid production and genes involved in lipid synthesis. Through transcriptomic analysis, 26,344 genes with varied expression were found. Functional enrichment analysis highlighted that pathways related to starch and sucrose metabolism, plant hormone signal transduction, and lipid accumulation were highly enriched among the differentially expressed genes. Additionally, expression variations of *SAD* and *FADs* among different varieties were explored. The analysis suggests that increased expression of genes such as *FAD3*, *FAD7*, and *FAD8* during the maturation stage may enhance the proportion of linolenic acid while reducing oleic acid content. This alteration could potentially stimulate fatty acid synthesis through a feedback regulation mechanism, thereby enhancing seed oil content. These findings provide a novel perspective on the feedback regulation mechanisms underlying lipid biosynthesis in *C. oleifera* seeds, offering significant insights to boost oil yield and improve tea oil quality.

## 1 Introduction

*Camellia oleifera* Abel., a tiny tree or shrub belonging to the Theaceae family, is a key source of woody oil in Asia, widely grown in countries such as China, the Philippines, Thailand, Japan, and South Korea (Ye et al., 2023). The *C. oleifera* seeds, known for their rich oil content, produce tea oil that is highly valued for its superior quality, significantly contributing to the economy. The fatty acids (FA) in tea oil predominantly consist of unsaturated types, essential for their high nutritional value (Zhang et al., 2022). Oleic acid, a monounsaturated FA, potentially plays a positive role in treating inflammation (Newton et al., 2022). Additionally, linoleic and linolenic acids in tea oil are crucial for lowering blood pressure (Tsukamoto and Sugawara, 2018). Tea oil is abundant in a wealth of bioactive elements like squalene, phytosterols, polyphenols, and tocopherols (Quan et al., 2022). By-products like Camellia seed cake, saponins, and fruit husks are utilized across various industries, including papermaking, chemical fiber, textiles, and pesticides (Liu et al., 2018). The cultivation of *C. oleifera* not only holds high economic value but also has a crucial role in soil conservation, and biodiversity maintenance (Gao et al., 2013; Zheng et al., 2008), thus promoting ecological stability and sustainable development.

Lipid biosynthesis in plant seeds primarily includes the de novo synthesis of FA in plastids and the synthesis of triacylglycerol in the endoplasmic reticulum (Bates et al., 2013). Acetyl-CoA carboxylase (*ACCase*), the initial enzyme in de novo FA synthesis, converts acetyl-CoA to malonyl-CoA and is widely regarded as the key rate-limiting step in this process (Sasaki and Nagano, 2004). After Malonyl-CoA: acyl carrier protein transacylase (*MCAT*) converts malonyl-CoA to malonyl-ACP, malonyl-ACP can further undergo a series of reactions catalyzed by fatty acid synthase (*FAS*), including condensation, reduction, dehydration, and re-reduction, thereby increasing the chain length. After seven cycles, palmitoyl-ACP is formed. Ketoacyl-ACP synthase II (*KASII*) further converts palmitoyl-ACP to stearoyl-ACP (Wei et al., 2012). Additionally, saturated acyl-ACP in plastids can be converted to unsaturated acyl-ACP under the desaturation effect of enzymes such as stearoyl-ACP desaturase (*SAD*), fatty acid desaturase 6 (*FAD6*), fatty acid desaturase 7 (*FAD7*), and fatty acid desaturase 8 (*FAD8*) (Kazaz et al., 2021). Fatty acyl-ACP thioesterase (*FAT*) is responsible for hydrolyzing acyl-ACP, ending the de novo synthesis of FA, and releasing free fatty acids (FFA). Different types of FAT have specific substrate selectivity. Fatty acyl-ACP thioesterase A (*FATA*) mainly hydrolyzes unsaturated long-chain acyl-ACP, while fatty acyl-ACP thioesterase B (*FATB*) tends to hydrolyze saturated long-chain acyl-ACP (Jing et al., 2011). The released FFA are converted to acyl-CoA by long-chain acyl-CoA synthetase (*LACS*) and participate in the assembly of triacylglycerol (TAG) in the endoplasmic reticulum (Bai et al., 2022). Glycerol-3-phosphate acyltransferase (*GPAT*), 1-acylglycerol-3-phosphate acyltransferase (*LPAAT*), phosphatidic acid phosphatase (PAP), and diacylglycerol acyltransferase (*DGAT*) catalyze four enzyme-catalyzed reactions in the Kennedy pathway, which produces TAG from glycerol 3-phosphate (G3P) and acyl-CoAs (Maraschin et al., 2019). In addition to TAG synthesis through the Kennedy pathway, recent discoveries have revealed more complex pathways centered around phosphatidylcholine (Bates and Browse, 2012; Chapman and Ohlrogge, 2012).

In recent years, rising demand for healthy foods and premium edible oils has spotlighted the FA composition and content in *C. oleifera* seeds as a key research area. Consequently, unraveling the biosynthetic mechanisms of *C. oleifera* FAs holds crucial practical value for enhancing tea oil quality to meet market demands. Advancements in omics technologies have spurred numerous studies into the biosynthesis and metabolism of FAs in C. oleifera seeds. Zhang Fanhang et al. (Zhang et al., 2021) revealed the developmental differences of Camellia oleifera seeds between different varieties through comparative transcriptome analysis of fruits from two Camellia oleifera varieties. Gong Wenfang et al. (Gong et al., 2020) found that the *WRINKLED1* (*WRI1*) transcription factor interacts with 17 key genes for lipid biosynthesis. Lin Ping et al. (Lin et al., 2018) proposed that in the later stages of seed development, high SAD activity and low FAD2 activity may contribute to an increase in oleic acid levels. Yang Jihong et al. (Yang et al., 2022) functionally characterized genes such as *SAD*, *FAD2*, *FAD3*, *DGAT1*, and *DGAT2*. Such research is pivotal for elucidating C. oleifera FA composition and metabolic pathways, thereby optimizing tea oil quality and boosting yields.

Given the complexity of the ploidy levels in *C. oleifera* and the abundance of varietal resources, extensive relevant studies are still necessary to comprehend the biosynthesis process of tea oil and the patterns of lipid accumulation. During plant lipid biosynthesis, FA manifests as both free and bound forms. Notably, the dynamic shifts in FFA during plant lipid synthesis have received limited focus. FFA, serving as intermediates between the FA de novo synthesis and TAG assembly pathways, are also influenced by processes like FA degradation. However, observing the changes in their content can assist in identifying potential feedback regulation within the lipid synthesis process. In this work, through transcriptome analysis of the seeds from three varieties of *C. oleifera* during the maturation stage, key genes in the biosynthesis process of tea oil were identified. Furthermore, by analyzing the composition ratio of FA and the dynamic changes in FFA content, this study delved into the lipid accumulation patterns in seeds of different *C. oleifera* varieties during the maturation stage. The findings in this research advance our comprehension of tea oil biosynthesis, offering vital insights to boost *C. oleifera* yields and enhance breeding quality.

## 2 Materials and methods

### 2.1 Plant materials

The National Engineering Research Center for Oil-Tea Camellia (Changsha, China), which is located at latitude 28°06′ N and longitude 113°01′ E, provided the experimental materials for this study. The base has an altitude range of 80–100 meters and is situated in an area with four different seasons, typical of a subtropical monsoon climate. Three eight-year-old *C. oleifera* varieties were selected in this study, namely XL192, XL108, and HH2. For clear and concise descriptions in the text, they will be referred to as L (XL192), M (XL108), and H (HH2), respectively. From September 14, 2022, the fruits on the Camellia oleifera trees were sampled every 14 days. The specific sampling dates were September 14 (T1), September 28 (T2), October 12 (T3) and October 26 (T4). Six fruits were selected from different positions of the tree each time, of which three fruits with a full appearance and uniform growth were selected as biological replicates. These samples were moved to the laboratory and stored at -80°C after being instantly frozen in liquid nitrogen.

### 2.2 Measurement of seed oil content

After removing the shells from the *C. oleifera* seeds, the kernels were dried in an oven at 60 ℃ until they reached a consistent weight. After drying, the kernels were processed in a grinder into a powder. A Soxhlet extractor was used to hold a filter paper bag containing 5g of the powder (m0), which had been precisely weighed. Petroleum ether was poured into two-thirds of the oil cup after its weight (m1) was measured. The water bath temperature was set to 70 °C, and the sample was reflux extracted for 6 hours. The weight of the oil cup (m2) was measured once the petroleum ether had evaporated to a consistent weight in the receiving bottle. The following formula was used to determine the amount of seed oil present: seed oil content (%) = (m2 - m1) / m0 × 100 In this case, m0 represents the sample weight (g), m1 the oil cup weight (g), and m2 the combined weight of the oil and oil cup (g).

### 2.3 Measurement of fatty acid content proportion

The FA composition was determined using a gas chromatograph, with each sample containing three biological replicates. A potassium hydroxide-methanol mixed solution (1.31 g potassium hydroxide, 10 ml methanol) was prepared. 4 milliliters of n-heptane and 60 microliters of oil were combined in a centrifuge tube and carefully stirred. Then, 200 milliliters of a potassium hydroxide-methanol solution were added, agitated for 30 seconds, and allowed to stand for approximately 30 minutes. 1 gram of sodium bisulfate was added to the centrifuge tube, shaken quickly, and allowed to settle for 20 minutes after standing. After the solution was clarified, the supernatant was drawn up with a syringe and added to the detection bottle. The detection bottles were loaded into the gas chromatograph in batches, and the FA content was quantitatively determined by the peak area of the chromatogram. The proportion of a certain FA was determined by comparing the peak area of the FA being analyzed to the total peak areas of all fatty acid components.

The determination parameters of the gas chromatograph are as follows: The flame ionization detector is set at 250 °C; the temperature of the sample injection port is 250 °C; the dimensions of the chromatographic column are 60 cm × 0.25 mm × 0.2 mm; the carrier gas is nitrogen; the split ratio is 1:50; the sample injection volume is 1 mL; the heating program is set to 50 °C (held for 2 minutes), 170 °C (rise by 10 °C/minute, held for 10 minutes), 180 °C (rise by 2°C/minute, held for 10 minutes), and 220 °C (rise by 4°C/minute, held for 22 minutes).

### 2.4 Measurement of free fatty acid content

After thawing the seed samples, mix 0.05 g with 150 µL methanol, 200 µL methyl tert-butyl ether, and 50 µL 36% phosphoric acid/water (which has been chilled to - 20°C). Vortex the mixture vigorously for 3 minutes at a speed of 2500 revolutions per minute (r/min), followed by centrifugation at 12 000 r/min for 5 minutes at a temperature of 4°C. Transfer 200 microliters of the clear liquid (supernatant) into a fresh centrifuge tube, use a nitrogen blower to dry it completely, and then introduce 300 microliters of a 15% boron trifluoride methanol solution. Vortex the mixture for 3 minutes at a speed of 2500 revolutions per minute, and subsequently incubate it in an oven set at 60°C for 30 minutes. Once the mixture has returned to ambient temperature, carefully add 500 microliters of n-hexane and 200 microliters of a saturated sodium chloride solution. After mixing, the mixture underwent centrifugation at 12,000 revolutions per minute for 5 minutes at a temperature of 4°C. Subsequently, 100 microliters of the upper n-hexane phase were collected for subsequent analysis via gas chromatography-mass spectrometry (GC-MS).

Derivatized samples were analyzed using a GC-EI-MS/MS system (GC, Agilent 7890B; MS, 7000D System). GC conditions: column, DB-5MS capillary column (30 m×0.25 mm×0.25 µm, Agilent); Carrier gas, high purity helium (purity >99.999%); The heating procedure was started at 40 ℃ (2 min), 30℃/min increased to 200℃ (1 min), 10℃/min increased to 240 ℃ (1 min), 5℃/min increased to 285℃ (3 min); traffic: 1.0 mL/min; inlet temperature: 230 ℃; injection volume: 1.0 µL. EI-MS/MS settings: Agilent 7890B-7000D GC-MS/MS System, Temperature, 230 °C; Ionization voltage, 70 eV; Transmission line temperature, 240 ℃; Four-stage rod temperature, 150 ℃; Solvent delay, 4 min; Scanning mode, SIM.

### 2.5 Analysis of RNA-Seq and Bioinformatics Exploration

TRIzol^TM^ Reagent (Invitrogen, CA, USA) was used to extract total RNA from 36 seed samples of three *C. oleifera* cultivars at four developmental phases. The purity of RNA was evaluated utilizing a NanoDrop spectrophotometer (Thermo Scientific, DE, USA), while its integrity was verified through an Agilent 2100 (Agilent Technologies, CA, USA). Finally, RNA degradation was assessed by conducting agarose gel electrophoresis (1%). RNA sequencing was performed by Beijing Allwegene Technology Company Limited (Beijing, China).

For each sample, 1.5 μg of RNA was utilized to enrich for mRNA using Oligo (dT) beads. 36 sequencing libraries were obtained using the NEBNext® UltraTM RNA Library Prep Kit for Illumina® (New England Biolabs, MA, USA), followed by PE 150 sequencing on the Illumina Novaseq 6000. Clean reads were generated by filtering out sequences from the raw data that contained sequencing adapters, had an N (unknown bases) ratio exceeding 10%, or were of low quality. The filtered clean reads were subsequently mapped to the *C. oleifera* reference genome (Lin et al., 2022) using STAR (v2.5.2b). Genes were considered differentially expressed if they had a corrected P-value of less than 0.005 and an absolute log2 (fold change) greater than 1.

### 2.6 Weighted gene correlation network analysis (WGCNA)

To study gene co-expression patterns, this study employed the WGCNA R package (version 1.72-5). The network creation and module identification were accomplished using default settings, with specific parameters as follows: an unsigned topological overlap matrix (TOM) was used for network construction; a soft threshold β (power β) was set to 8 to enhance the interconnectivity between modules; each module contained at least 100 genes; the branch merge cut height for module merging was set at 0.25 (i.e., merging co-expression modules with at least 75% similarity). Module eigengene (ME), representing the first principal component of all gene expression data in a module, reflected the overall expression pattern of that module and was used to explore associations with oil amounts. Module membership (MM) was identified by matching a gene’s expression profile with the ME of its appropriate module and was significantly linked to the module’s internal connection (K.in). Utilizing the MM and K.in, Cytoscape (Version 3.10.1) was used to construct regulatory networks of genes within the module.

### 2.7 Real-time quantitative PCR (qRT-PCR) verification

Ten genes relevant to lipid synthesis were chosen at random for quantitative real-time experiments to verify the data from RNA-seq. The *tubulin* gene was employed as an internal reference. Primer Premier 5 (Premier Biosoft, Palo Alto, CA, USA) was used to build the primers required to perform qRT-PCR assays (Supplementary Table S1). Total RNA from *C. oleifera* seeds was extracted using the RN53-EASYspin Plus Polysaccharide Polyphenol/Complex Plant RNA Rapid Extraction Kit (Aidlab, Beijing, China), and first-strand cDNA was obtained using the HiScript III 1st Strand cDNA Synthesis Kit (+gDNA wiper) (Vazyme, Nanjing, China). The cDNA was then diluted fivefold and used as the template for qRT-PCR, with each sample including three technical replicates. The 2^−ΔΔCt^ method was used to calculate the relative transcription levels of the genes.

## 3 Results

### 3.1 Total oil content and proportions of major FA components in seeds of different varieties

The results showed the H variety had a considerably higher oil content, exceeding that of the L and M varieties by 34.97% and 34.30%, respectively (Fig. 1). While the M variety exhibited a marginally higher oil content than the L variety, this difference was not statistically significant. Oleic acid emerged as the dominant FA component in all varieties’ kernels, making up over 70% of the FA composition, with palmitic acid, stearic acid, linoleic acid, and linolenic acid following in sequence. Notably, an inverse relationship was observed between oil content and oleic acid proportion, with a corresponding increase in linoleic acid.

**Fig. 1.**
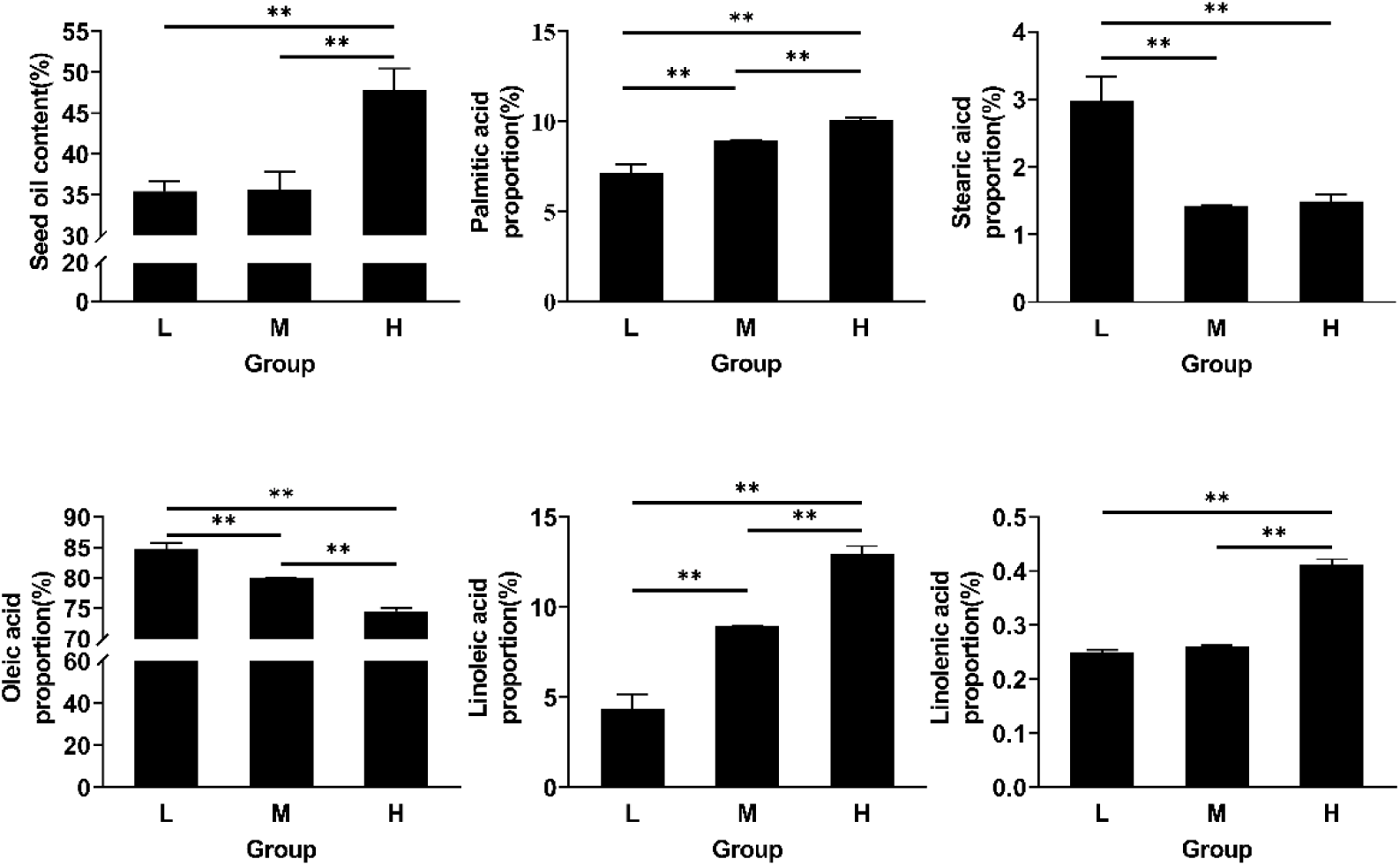
Total oil content and proportions of major FA components in seeds of three varieties (T4, October 26). The data were analyzed through one-way ANOVA and the Tukey test to identify significant differences (* indicates p ≤ 0.05 and ** indicates p ≤ 0.01). They’re shown as mean ± SD from three biological replicates. L, XL192; M, XL108; H, HH2.

Overall, significant variations in fatty acid (FA) composition were observed among the different *C. oleifera* varieties examined in this study. Specifically, in FA composition, the proportions of stearic acid and oleic acid were considerably greater in the L variety than in the M and H varieties. In contrast, the M variety showed a higher preference for oleic acid and linoleic acid proportions. H variety had significantly higher proportions of linoleic acid and linolenic acid than the M and L varieties, indicating significant differences in unsaturated FA preference among these varieties.

### 3.2 Dynamic changes of free fatty acid content in kernels of different varieties

This study further explored FA accumulation patterns across different varieties by analyzing FFA composition and content in kernels at four maturation stages. As shown in Fig. 2A, across all varieties, the contents of free palmitic acid, free stearic acid, free oleic acid, and free linoleic acid exhibited a general upward trend over time. Conversely, free linolenic acid content demonstrated a general decline throughout the period. It is noteworthy that at T2, the L and M varieties experienced a slight decrease in free linoleic acid content compared to T1. Furthermore, in the H variety, free linolenic acid content surged by 41.04% from T1 to T2, then plummeted by 60.05% from T2 to T3. During the T3∼T4 stage, the L variety saw significant increases in free stearic and oleic acid contents.

**Fig. 2.**
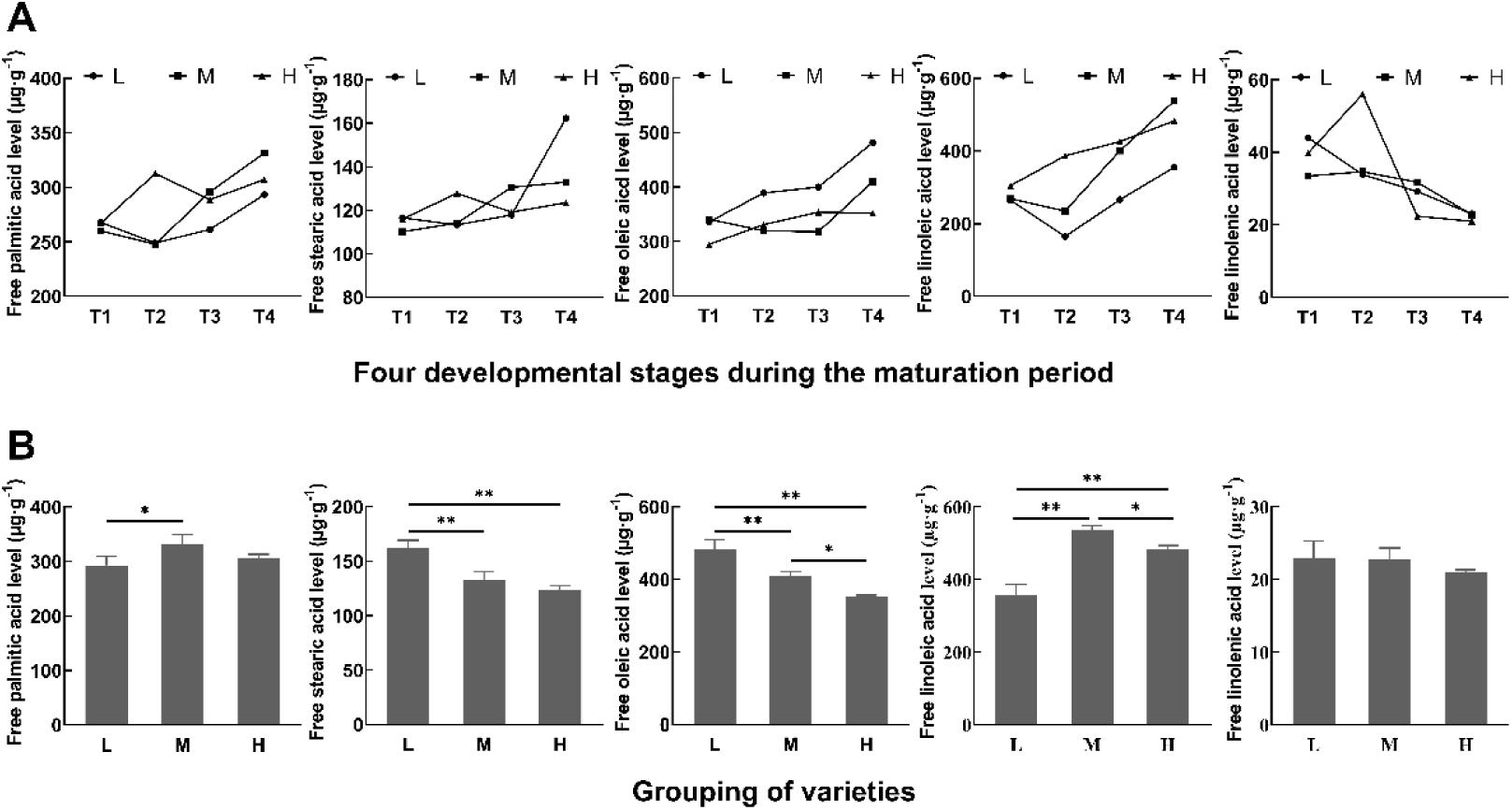
A: Temporal trends of major free fatty acids from T1 to T4. B: Levels of major free fatty acids in kernels of each variety at T4. The data were analyzed through one-way ANOVA and the Tukey test to identify significant differences (* indicates p ≤ 0.05 and ** indicates p ≤ 0.01). They’re shown as mean ± SD from three biological replicates. L, XL192; M, XL108; H, HH2; T1, September 14; T2, September 28; T3, October 12; T4, October 26.

As shown in Fig. 2B, at T4, alongside an increase in overall oil content, there was a decreasing trend in the contents of free stearic and oleic acids in the kernels. Interestingly, at T4, despite the linoleic acid proportion being significantly higher in the H variety compared to the L and M varieties (Fig. 1), the M variety exhibited the highest free linoleic acid content, followed by the H and L varieties (Fig. 2B). Moreover, at T4, the free linolenic acid content in the H variety was comparable to that in the L and M varieties, showing no significant differences (Fig. 2B). However, in terms of linolenic acid proportion, the H variety was much higher than the L and M varieties (Fig. 1). These observations indicate that at T4 in the H variety, a substantial portion of linoleic and linolenic acids may be present as bound FA.

### 3.3 Characterization of the transcriptomes and identification of differentially expressed genes (DEGs)

The molecular mechanisms of oil biosynthesis during *C. oleifera* seed development was investigated through RNA-seq analysis on samples from various developmental stages. Three biological replicates were established for each developmental stage of the seed samples to ensure data reliability. From the 36 libraries, approximately 1.571 billion high-quality clean reads were generated in total (Supplementary Table S2), which were used for subsequent alignment analysis. Of these, approximately 1.405 billion (89.45%) high-quality reads were successfully mapped, of which 1.235 billion (78.61%) reads were uniquely mapped (Supplementary Table S3).

Regarding gene expression, the number of valid expressed genes with FPKM values exceeding 1.0 in L variety at T1, T2, T3, and T4 stages was 28781, 26870, 27151, and 24384, respectively; the corresponding numbers for M variety were 26431, 25573, 25582, and 25374; and those for H variety were 25062, 28617, 24779, and 24,833 (Supplementary Table S4). Generally, the count of validly expressed genes in the L and M varieties gradually decreased through the ripening process, while in the H variety, it demonstrated an initial rise followed by a decline.

In the pairwise comparisons between varieties at identical stages, 36244 DEGs were identified between L and M varieties (L1vsM1, L2vsM2, L3vsM3, and L4vsM4) (Fig. 3A), while 36918 DEGs were identified between H and M varieties (H1vsM1, H2vsM2, H3vsM3, and H4vsM4) (Fig. 3B). Furthermore, more DEGs were identified in the pairwise comparisons of adjacent stages within the variety with higher oil content (Fig. 3F). Specifically, 24031 DEGs were identified in L variety (L2vsL1, L3vsL2, and L4vsL3) (Fig. 3C), 29815 DEGs were identified in M variety (M2vsM1, M3vsM2, and M4vsM3) (Fig. 3D), and the most DEGs were identified in H variety (H2vsH1, H3vsH2, and H4vsH3), with a total of 33505 DEGs (Fig. 3E).

**Fig. 3.**
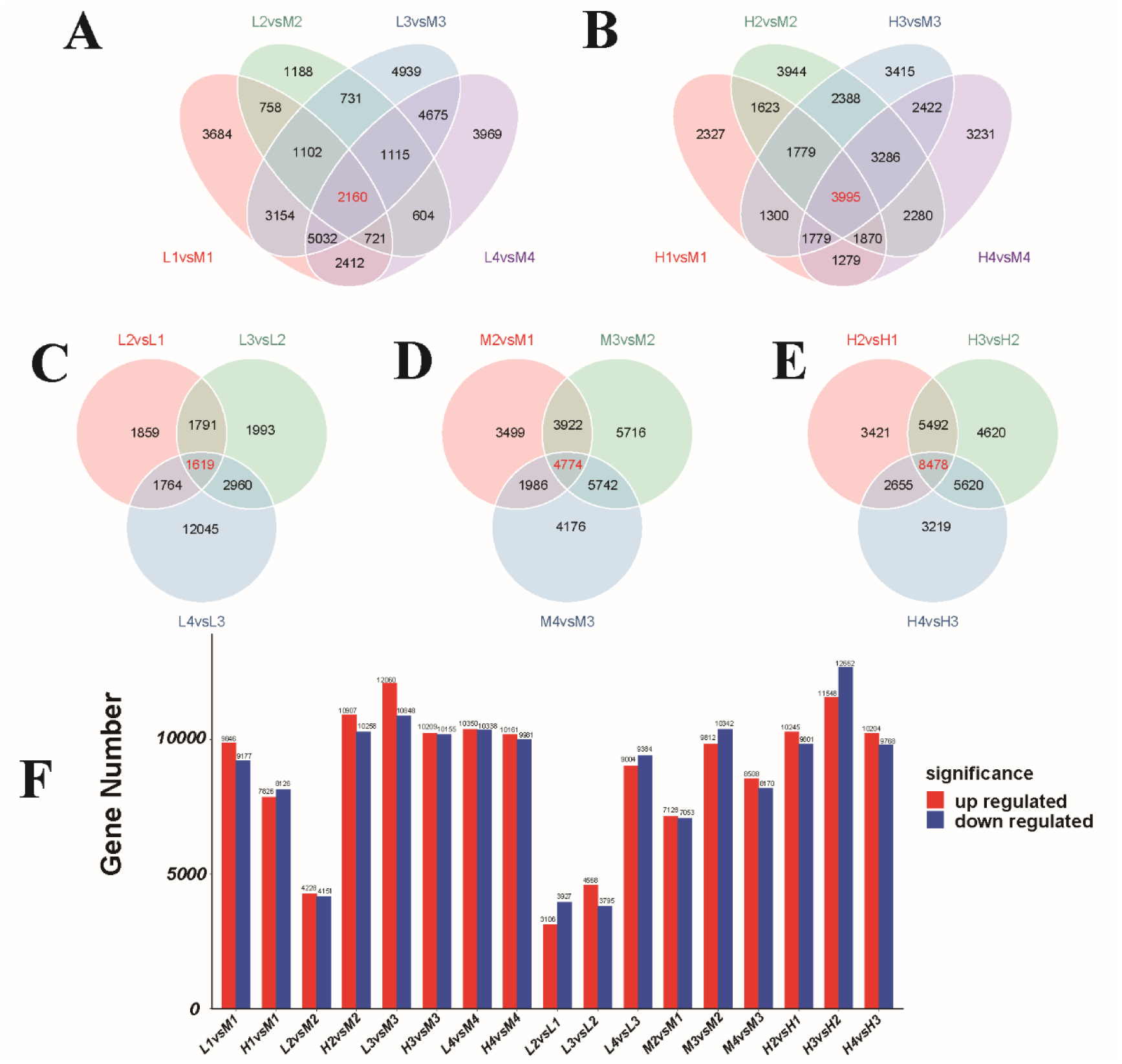
A: Venn diagram of DEGs between L and M varieties. B: Venn diagram of DEGs between H and M varieties. C: Venn diagram of DEGs between adjacent stages of L variety. D: Venn diagram of DEGs between adjacent stages of the M variety. E: Venn diagram of DEGs between adjacent stages of the H variety. F: Number of DEGs in different pairwise comparisons. Red represents up-regulated genes, and blue represents down-regulated genes.

Across all same-stage pairwise comparisons between L and M varieties and H and M varieties (L1vsM1, H1vsM1, L2vsM2, H2vsM2, L3vsM3, H3vsM3, L4vsM4, H4vsM4), a total of 27981 annotated DEGs were identified (Supplementary Table S5).

At the same time, 27600 DEGs were found in the pairwise comparisons of adjacent stages within L, M, and H varieties (L2vsL1, L3vsL2, L4vsL3, M2vsM1, M3vsM2, M4vsM3, H2vsH1, H3vsH2, H4vsH3) (Supplementary Table S5). A total of 26 344 DEGs, common across both comparison categories, were designated as significant differentially expressed genes (SigDEGs). These SigDEGs were identified in both same-stage pairwise comparisons and adjacent-stage pairwise comparisons (Supplementary Table S5), highlighting their substantial value in exploring gene expression dynamics.

The identified SigDEGs underwent further analysis through GO and KEGG enrichment (Fig. 4). GO enrichment analysis revealed that, within the biological process (BP) category, the predominant GO terms included metabolic process, cellular metabolic process, and organonitrogen compound metabolic process. For the cellular component (CC) category, leading GO terms comprised membrane, intracellular, and organelle. In the molecular function (MF) category, the top three GO terms were catalytic activity, small molecule binding, and nucleotide binding (Fig. 4A). Moreover, KEGG enrichment analysis identified 51 significantly enriched pathways, encompassing plant hormone signal transduction, starch and sucrose metabolism, photosynthesis, fatty acid elongation, steroid biosynthesis, ABC transporters, arachidonic acid metabolism, pantothenate and CoA biosynthesis, among others (Fig. 4B).

**Fig. 4.**
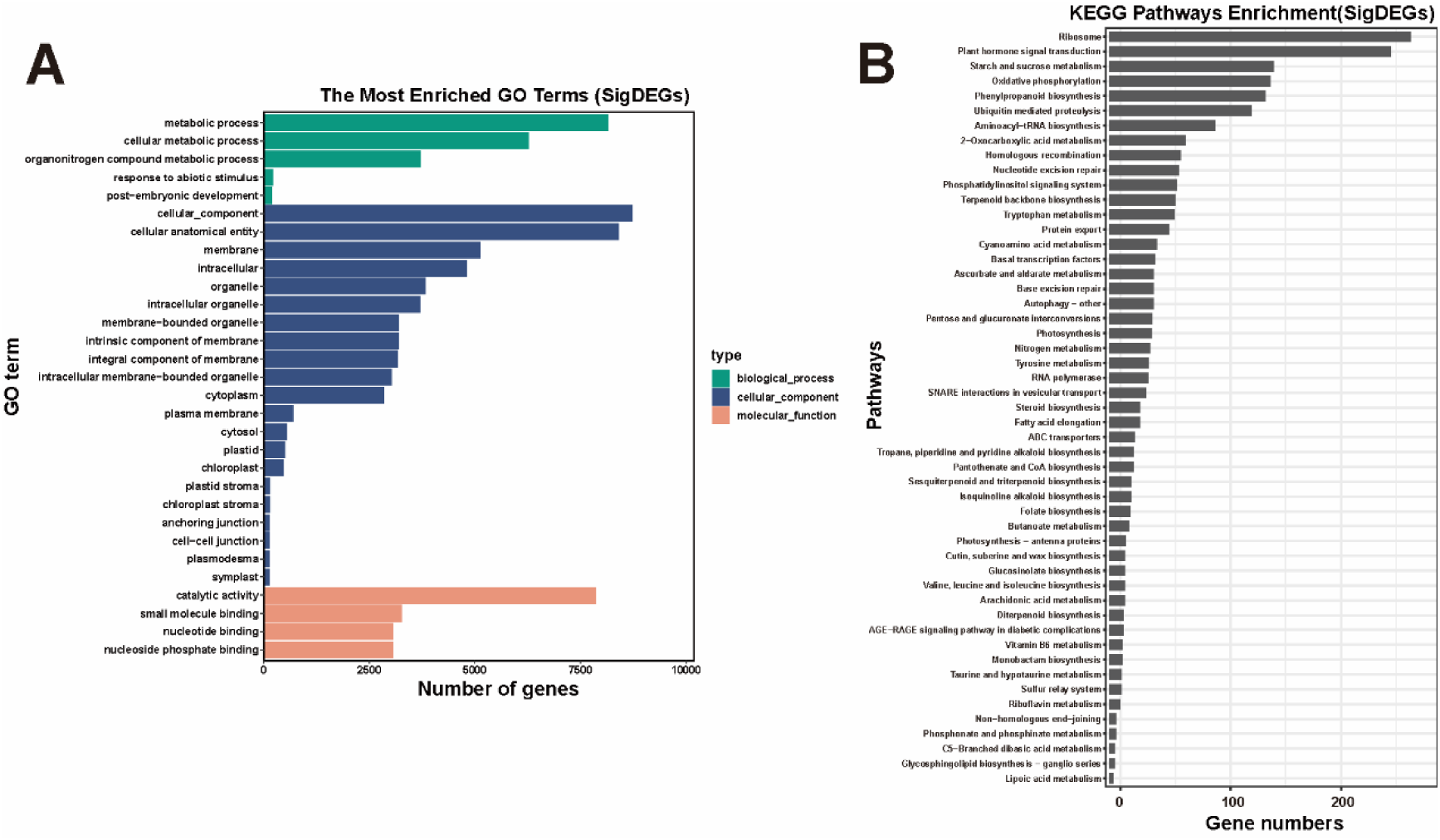
GO and KEGG enrichment analysis of SigDEGs. A: GO enrichment analysis of SigDEGs. B: KEGG enrichment analysis of SigDEGs

### 3.4 Weighted gene co-expression network analysis of SigDEGs

This study employed the median absolute deviation (MAD) to filter out noise among SigDEGs, selecting 14,608 SigDEGs with MAD values ≥ 1.0 for WGCNA analysis. The analysis divided these 14,608 SigDEGs into 12 co-expression modules (Fig. 5A), of which the blue module (4943 genes) was most correlated with linoleic acid, while the green-yellow (435 genes) and dark-green modules (4232 genes) both showed high correlations with linolenic acid (Fig. 5B). Despite the green-yellow module showing the highest correlation with linolenic acid, it contained only a few genes involved in lipid syntheses, such as FATB, GPAT, and LPCAT. In contrast, large numbers of lipid synthesis-related genes were found in the blue and dark-green modules, including *ACCase*, *KAR*, *KASI*, *KASII*, *KASIII*, *SAD*, *FAD6*, *FAD7*, *FAD8*, *FATB*, *LACS*, *GPAT*, and *PDAT* genes (Supplementary Table S6).

**Fig 5.**
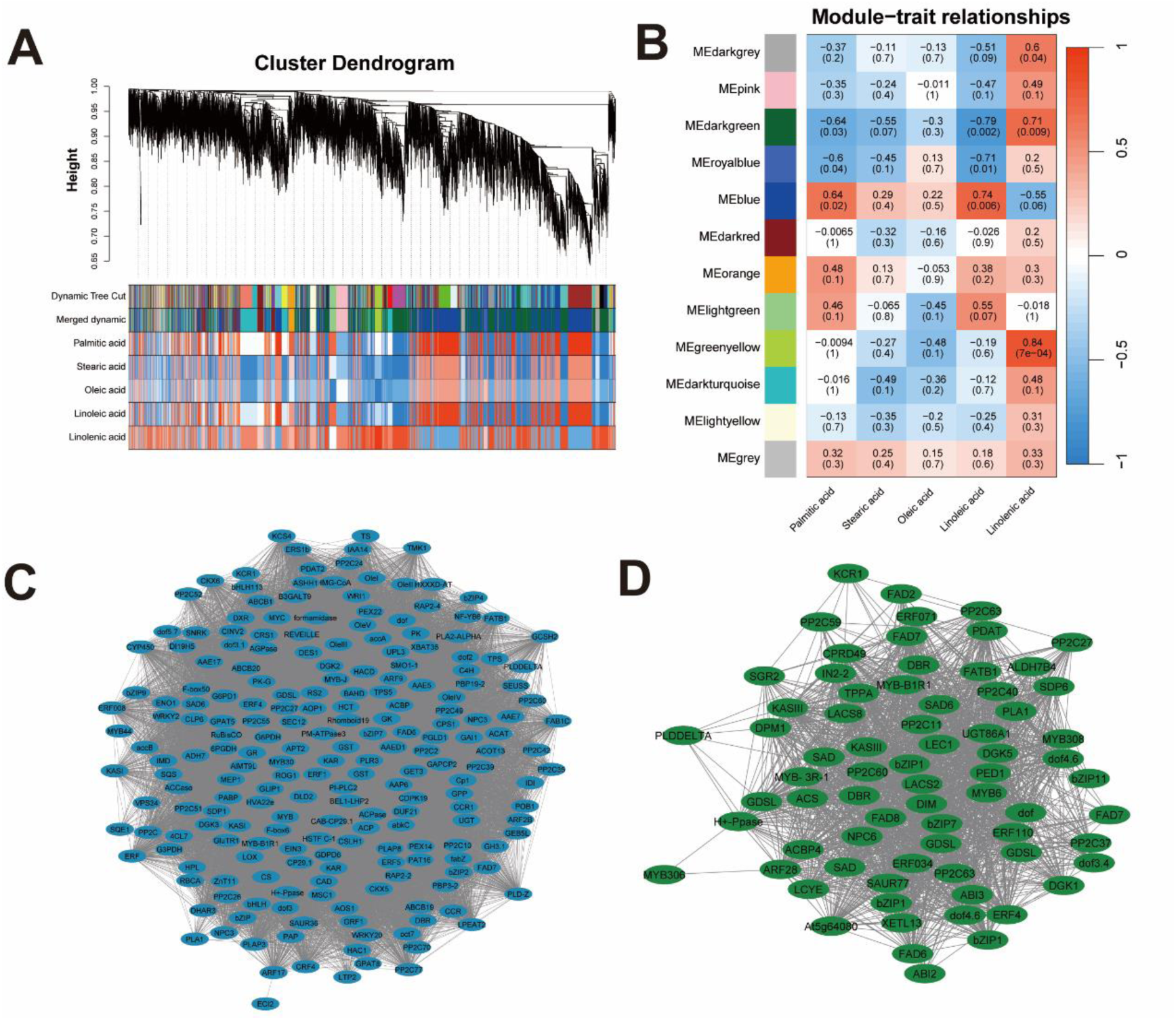
A: Clustering dendrograms of SigDEGs. B: Heatmap showing the correlation between modules and free fatty acid content. C: Network relationship of genes related to oil synthesis and transcription factors in the blue module. D: Network relationship of genes related to oil synthesis and transcription factors in the dark-green module.

Notably, both modules contained *FAD6* and *FAD7*, while *FAD8* only appeared in the Dark-green module (Fig. 5C, Fig. 5D). Furthermore, transcription factors (TFs) identified in these modules underscore their crucial regulatory roles in FA synthesis. The TFs identified in the two modules included *MYB*, *bZIP*, *DOF*, *WRI1*, *LEC*, *ABI3*, *WRKY*, *bHLH*, *GRF*, *ARF*, and *ERF*. Protein kinases related to abscisic acid signaling, such as *PP2C* and *SNRK*, were also found in the two modules. This finding highlights the significant roles of phytohormones, including ethylene, auxin, and abscisic acid, in lipid synthesis.

### 3.5 Expression patterns of genes related to oil synthesis pathways during the maturation of kernels of different varieties

Within the de novo FA synthesis pathway, genes such as *ACCase*, *KAS I*, *KAS II*, *KAS III*, *KAR*, *HAD*, *EAR*, and *SAD* in the H variety were significantly upregulated at T4 (Fig. 6), indicating that FA synthesis activity may be relatively more active in the T3∼T4 stage of the H variety. Specifically, *FAD6* experienced significant upregulation during the T2∼T3 stage in the H variety. Notably, the three transcripts of *FAD7* in the H variety were mainly upregulated in the T1∼T2 and T2∼T3 stages, while *FAD8* was upregulated in the T3∼T4 stage. In comparison, the *FAD7* gene in the M variety was mainly upregulated in the T2∼T3 stage, but *FAD8* gene expression levels were almost unchanged across stages. In the L variety, *FAD7* and *FAD8* only showed slight upregulation in the T2∼T3 stage.

**Fig. 6.**
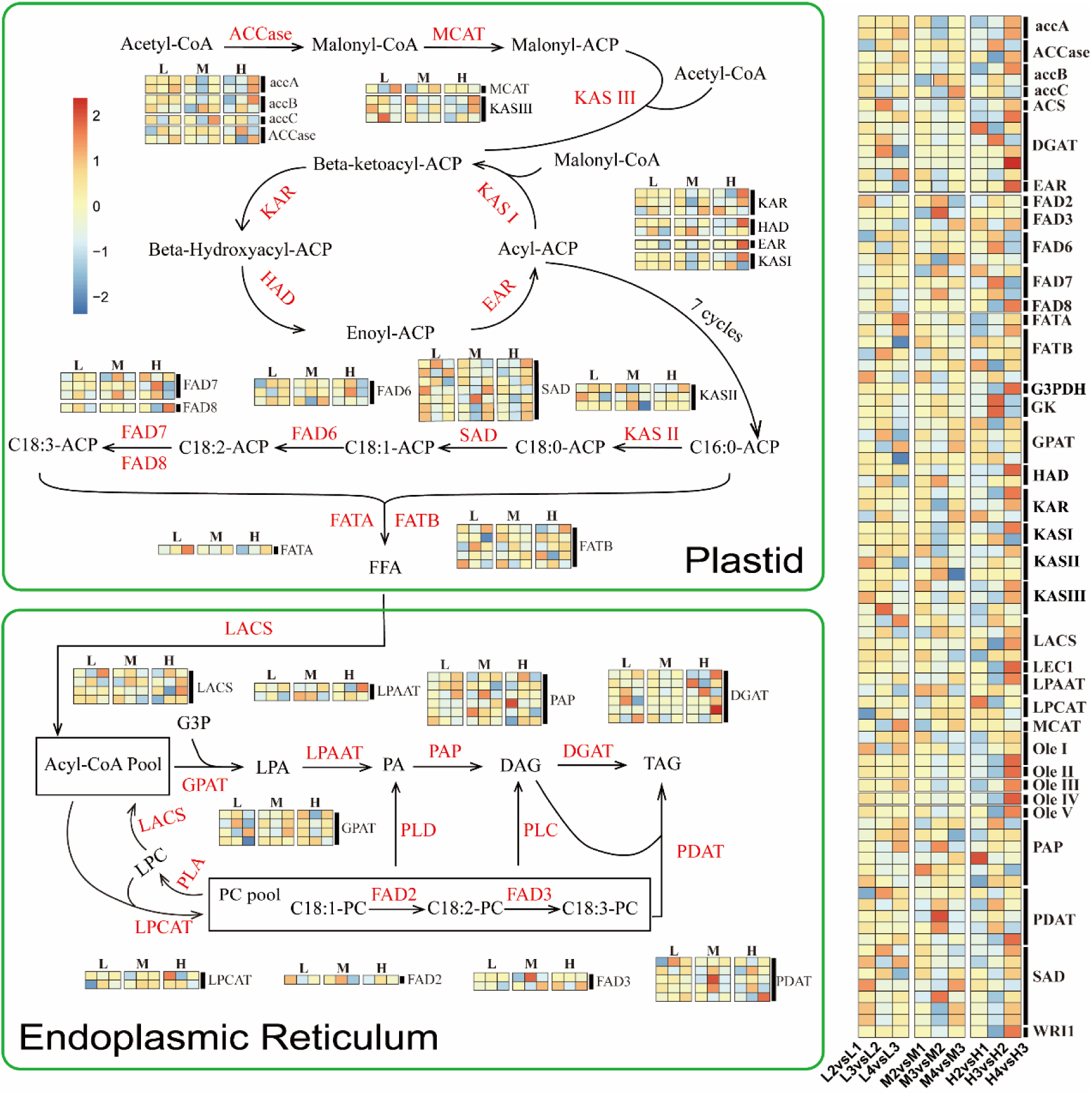
Oil synthesis pathway map and heatmap of gene expression related to oil synthesis. Heatmaps were generated using log2FC values of gene expression between adjacent stages of each variety (Supplementary Table S7). Reaction abbreviations: *accA*, acetyl-coenzyme A carboxylase carboxyl transferase subunit alpha; *accB*, biotin carboxyl carrier protein of acetyl-CoA carboxylas; *ACCase*, Acetyl-CoA carboxylase; *ACS*, acetyl-coenzyme A synthetase; *DGAT*, diacylglycerol acyltransferase; *EAR*, enoyl-[acyl-carrier-protein] reductase; *FAD2*, fatty acid desaturase 2; *FAD3*, fatty acid desaturase 3; *FAD6*, fatty acid desaturase 6; *FAD7*, fatty acid desaturase 7; *FAD8*, fatty acid desaturase 8; *FATA*, acyl acyl-carrier-protein thioesterase type A; *FATB*, acyl acyl-carrier-protein thioesterase type B; *G3PDH*, glycerol-3-phosphate dehydrogenase; *GK*, glycerol kinase; *GPAT*, sn-glycerol-3-phosphate acyltransferase; *HAD*, 3-hydroxyacyl-[acyl-carrier-protein] dehydratase; *KAR*, ketoacyl-ACP reductase; *KASI*, ketoacyl-ACP synthase I; *KASII*, ketoacyl-ACP synthase II; *KASIII*, ketoacyl-ACP synthase III; *LACS*, long chain acyl-CoA synthetase; *LEC1*, leafy cotyledon 1; *LPAAT*, lysophosphatidic Acid Acyltransferase; *LPCAT*, 1-acylglycerol-3phosphate acyltransferase; *MCAT*, malonyl-CoA-acyl carrier protein transacylase; *Ole I*, oleosin I; *Ole II*, oleosin II, *Ole III*, oleosin III; *Ole IV*, *ole*osin IV; *Ole V*, oleosin V; *PAP*, Phosphatidate Phosphatase; *PDAT*, Phospholipid: Diacylglycerol Acyltransferase; *PLA*, Phospholipase A; *PLC*, Phospholipase C; *PLD*, Phospholipase D; *SAD*, stearoyl-ACP desaturase; *WRI1*, wrinkled1.

**Fig. 7.**
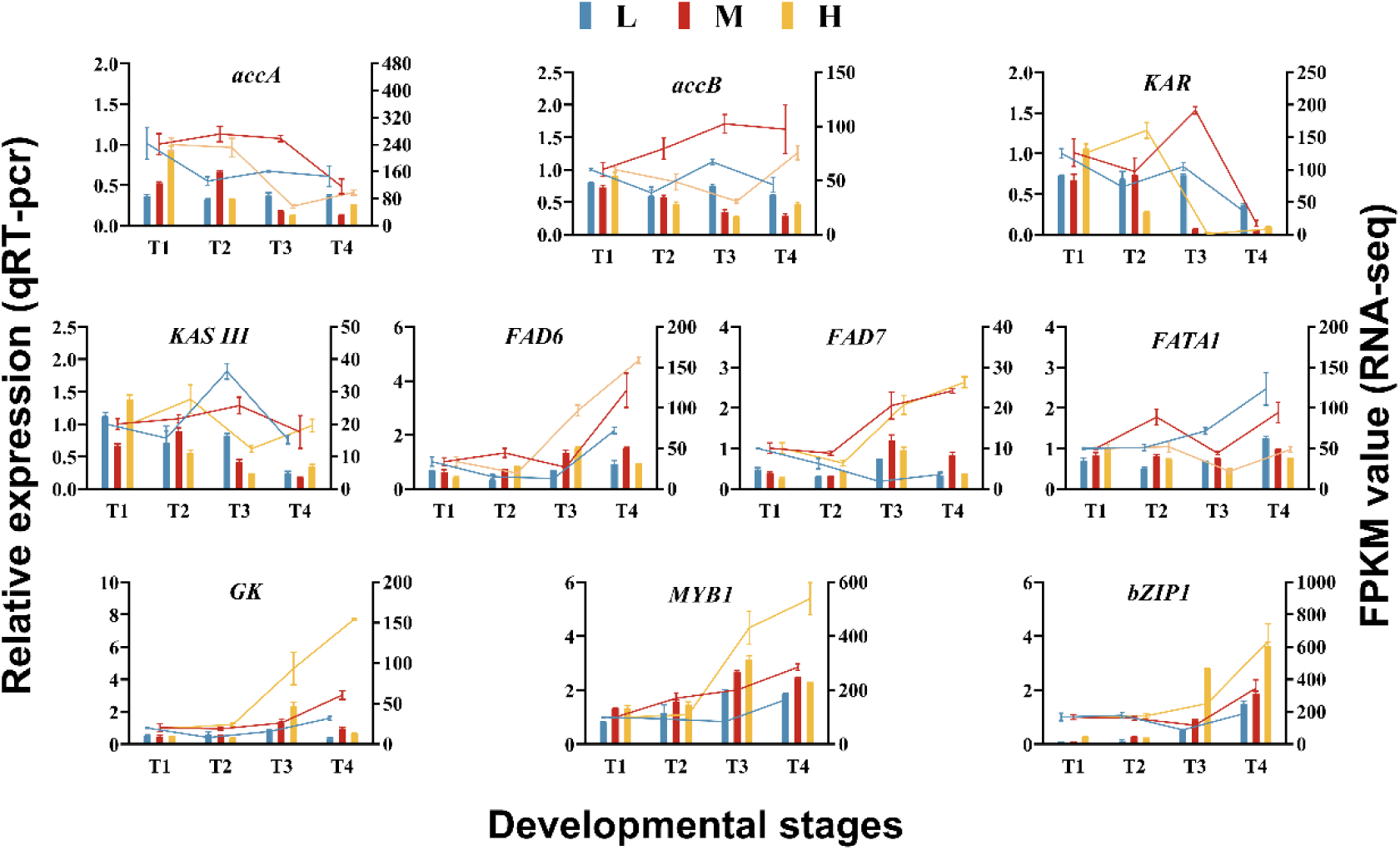
Comparison and validation of qRT-PCR and RNA-seq results. The bar chart shows the expression level of FPKM (right Y-axis), and the line chart shows the relative expression level of qRT-PCR (left Y-axis). The error bars in the figure represent the standard deviation (SD). The abbreviations and full names of the related genes are as follows: *accA*, acetyl-coenzyme A carboxylase carboxyl transferase subunit alpha; *accB*, biotin carboxyl carrier protein of acetyl-CoA carboxylase; *KAR*, ketoacyl-ACP reductase; *KASIII*, ketoacyl-ACP synthase III; *FAD6*, fatty acid desaturase 6; *FAD7*, fatty acid desaturase 7; *FATA1*, acyl acyl-carrier-protein thioesterase type A1; *GK*, glycerol kinase; *MYB1*, myeloblastosis transcription factor 1; *bZIP1*, basic leucine zipper 1.

Within the TAG synthesis pathway, the *FAD3* gene in the H variety was observed to be upregulated at both the T1∼T2 and T3∼T4 stages, up-regulated in the M variety at the T2∼T3 stage, and showed no significant changes in all stages in the L variety. The difference in *FAD* expression between different varieties may explain the difference in linolenic acid content. In addition, compared to the L variety and M variety, the corresponding transcripts of the *DGAT* gene in the H variety were up-regulated in all stages, indicating that TAG assembly activity in the H variety was also relatively active in all stages. Particularly at the T3∼T4 stage, significant upregulation was observed for the *LPAAT*, *LACS*, *DGAT*, and *PDAT* genes in the H variety. Notably, the *Ole* gene in the H variety was significantly upregulated at the T3∼T4 stage, suggesting a pivotal role for oleosin in lipid synthesis. *G3PDH* provides the glycerol backbone for TAG biosynthesis, and its high expression at the T3∼T4 stage provides sufficient raw materials for TAG biosynthesis. Furthermore, the significant upregulation of *WRI1* and *LEC1* in the H variety at the T3∼T4 stage underscores their critical roles in regulating lipid synthesis pathways.

In conclusion, during the T3∼T4 stage, multiple genes related to lipid synthesis in the H variety were co-upregulated, which may be the main reason why its oil content is higher than that of the L variety and the M variety. The enzymes encoded by genes such as *FAD3*, *FAD7*, and *FAD8* are involved in the synthesis of linolenic acid. The high expression of these genes in the H variety effectively increased the content of linolenic acid in the seeds.

### 3.6 Verification of RNA-seq data with RT-qPCR

Ten DEGs associated with lipid synthesis were picked at random for qRT-PCR experiments to confirm the veracity of the RNA-seq data. The outcomes revealed that the expression patterns of the majority of these genes were closely aligned between the qRT-PCR results and the RNA-seq data. This high level of similarity confirms the reliability of the RNA-seq data, underscoring its value in identifying and analyzing gene expression related to lipid production.

## 4 Discussion

In this work, it was noticed that the oil content of the three *C. oleifera* varieties on October 26 (T4) decreased in the order of HH2 (H), XL108 (M), and XL192 (L). Notably, the L variety exhibited a significantly higher stearic acid content compared to the M and H varieties. An intriguing inverse relationship emerged between the oil content and oleic acid proportion, while a direct relationship was observed with the proportion of linoleic acid and linolenic acid. Transcriptome sequencing of the seeds identified 43 865 DEGs, among which 26 344 genes were commonly identified across both the pairwise comparisons between varieties within the same period (L1vsM1, H1vsM1, L2vsM2, H2vsM2, L3vsM3, H3vsM3, L4vsM4, H4vsM4) and the pairwise comparisons between adjacent stages within each group (L2vsL1, L3vsL2, L4vsL3, M2vsM1, M3vsM2, M4vsM3, H2vsH1, H3vsH2, H4vsH3). In-depth mining of these 26,344 DEGs uncovered significant varietal differences in key genes, including *SAD* and *FAD*. Additionally, TFs and phytohormones were identified as playing crucial roles in FA biosynthesis in the kernels. Overall, the insights gained from this research offer valuable guidance and constitute an important reference for enhancing the breeding quality of *C. oleifera*.

### 4.1 The coordinated expression of *SAD* and *FAD* genes affects the FA composition in C. oleifera Kernels

Previous research has identified that tea oil kernels have two peaks of oil accumulation, with the second peak occurring from mid-September to late October (Zhang et al., 2021). In this work, the dynamic changes of FFA revealed differences in lipid accumulation among the three varieties during the second peak of oil accumulation. These differences predominantly manifested themselves in the varieties’ preference for FA unsaturation levels. The L variety is inclined to produce FAs with lower desaturation levels, notably stearic and oleic acids. In contrast, the M variety tends to produce FA with a medium degree of desaturation, like oleic acid and linoleic acid. The H variety tends to produce highly unsaturated FA, especially linoleic acid and linolenic acid. Previous studies have demonstrated that the elevated oleic acid level in tea oil kernels is associated with sustained high levels of transcription in the *SAD* gene and coordinated inadequate transcription of the *FAD2*, *FAD3*, *FAD7*, and *FAD8* genes (Lin et al., 2018; Wu et al., 2019; Yang et al., 2022).

During seed oil biosynthesis, *SAD*, *FAD6*, *FAD7*, and *FAD8* in plastids, alongside *FAD2* and *FAD3* in the endoplasmic reticulum, enhance FA desaturation by introducing double bonds (Bates et al., 2013). In plastids, stearic acid undergoes successive introduction of one double bond by SAD, FAD6, and FAD7/FAD8 to form oleic, linoleic, and linolenic acids, respectively (He et al., 2020). In the endoplasmic reticulum, oleic acid in phosphatidylcholine can be introduced with one double bond by *FAD2* and *FAD3* to generate linoleic acid and linolenic acid, respectively (Fig. 6). In this study, *SAD* gene in the L variety had corresponding transcript upregulation at different stages, while genes including *FAD3*, *FAD7*, and *FAD8* were expressed at low levels at most stages, limiting the conversion from oleic to linoleic and linolenic acids, leading to oleic acid accumulation. Such findings align with those from previous research. For the H variety, increased activity of FAD3, FAD7, and FAD8 suggested enhanced linolenic acid synthesis at all stages, consequently reducing the oleic acid proportion.

### 4.2 Potential feedback regulation in FA biosynthesis

Despite the H variety’s relatively low oleic acid proportion, this appears to enhance carbon source entry into the FA biosynthesis pathway. High expression of *ACCase* is a hallmark of carbon flux through the de novo FA biosynthesis pathway and plays a key role in boosting seed oil production (Wang et al., 2022). Additionally, *KASⅢ*initiates the first condensation process during the first cycle of FA synthesis, marking it as another vital rate-limiting enzyme (Nofiani et al., 2019). Thus, these genes’ co-expression is advantageous for kernel oil content enhancement. In this study, compared to the L and M varieties, nearly all genes associated with the de novo FA synthesis pathway of the H variety showed a synergistic up-regulation effect at the T3∼T4 stage, which is likely linked to the low level of oleic acid. Within biosynthetic pathways, downstream product accumulation can feedback-inhibit initial enzyme activity, thereby regulating the synthesis rate of the pathway and maintaining metabolic balance. Studies have shown that the accumulation of 18:1-ACP can inhibit the activity of *ACCase* in rapeseed, but does not affect the extension of FA (Andre et al., 2012). In this study, the transformation of oleic acid into linoleic acid was active at all stages in the H variety, causing a reduction in free oleic acid content from T3 to T4 (Fig. 2), which might facilitate the entry of carbon sources toward the FA biosynthesis pathway at the T4 stage through a potential feedback regulation mechanism.

Triacylglycerol usually remains in oil bodies after assembly, with *ole* being the most abundant protein on the phospholipid membrane (Frandsen et al., 2001; Shimada and Hara-Nishimura, 2010). Despite the ability of oil bodies to form without oleosin, numerous pieces of evidence imply that *ole* serves an essential part of preserving oil body stability (Wijesundera and Shen, 2014; Idogawa et al., 2018; Yee et al., 2021). In this research, multiple *ole* were significantly elevated within the H variety at the T3∼T4 stages, accompanied by a significant up-regulation of TAG biosynthesis genes such as *LPAAT*, *DGAT*, and *PDAT* (Fig. 6). *Ole1* overexpression has been shown to promote oil accumulation by inhibiting TAG degradation-associated transport proteins and lipases, and by activating enzymes related to TAG assembly (Zhang et al., 2019; Xu et al., 2020; Hu et al., 2023). TAG is one of the ultimate products for the Kennedy pathway. However, there is currently no evidence whether the content of “free TAG” can feedback-regulate the enzymes or transcription factors involved in TAG biosynthesis.

### 4.3 The regulatory roles of TFs in FA biosynthesis

The *WRI1* acts as a pivotal regulator within oilseed FA synthesis pathways, controlling multiple enzymes for glycolysis and de novo FA synthesis (Kong and Ma, 2018; Kong et al., 2019). Additionally, *LEC1* regulates FA synthesis depending on *FUS3*, *ABI*, and *WRI1* (Mu et al., 2008). Elevated levels of *WRI1* and *LEC1* are strongly connected to enhanced seed oil production (Tan et al., 2011; Huang et al., 2023). Thus, the simultaneous upregulation of *ACCase*, *EAR*, *HAD*, *KAR*, *KASI*, *KASII*, *KASIII*, and *SAD* genes in the H variety during T3∼T4 stages may correlate with elevated *LEC1* and *WRI1* gene expression. Furthermore, although previous studies have suggested that TAG biosynthesis in the endoplasmic reticulum is not regulated by *WRI1* (Maeo et al., 2009), recent research has indicated that WRI1 can target genes related to TAG biosynthesis, such as *G3PHD*, *DGAT2*, and *PDAT* (Kuczynski et al., 2022; Yuan et al., 2022). The data from this research indicate that *G3PHD*, *DGAT*, and *PDAT* also exhibited significant up-regulation in the H variety at the T3∼T4 stage, which further supports the potential regulatory role of *WRI1* in TAG biosynthesis.

Earlier research indicates that R2R3-type *MYB* transcription factors generally exert a negative influence on oil biosynthesis in the *C. oleifera* seeds, with a minority being implicated in long-chain FA synthesis (Gong et al., 2020; Li et al., 2022). Moreover, research has demonstrated that the overexpression of *DOF* stimulates oil buildup in many experiments. (Su et al., 2017; Wang et al., 2007; Jia et al., 2019). In this work, 9 *DOFs* were identified from the blue and dark-green modules based on WGCNA (Supplementary Table S6), which may be closely related to oil synthesis. The production of oil and the development of seeds are strongly influenced by *bZIP* transcription factors. In Arabidopsis (Mendes et al., 2013), *bZIP67* can increase the content of linolenic acid in Arabidopsis seeds by activating the desaturase FAD3. Further research shows that *bZIP67* can cooperate with *LEC1*, *AREB3*, and *ABI3* to oversee gene expression throughout the seed development process (Jo et al., 2020). In addition, transgenic Arabidopsis seedlings have also been shown to have higher lipid contents when GmbZIP123 is overexpressed (Song et al., 2013). This suggests that *bZIP* positively contributes to seed lipid accumulation. In this work, according to WGCNA, a total of 10 *bZIP*s that might be associated with oil biosynthesis were found (Supplementary Table S6).

### 4.4 Crucial role of phytohormones in FA production

Phytohormones are essential for the production of FA. (Shahid et al., 2019; Nguyen et al., 2021). According to this study’s WGCNA data, FA biosynthesis is significantly impacted by *ARF*, *ERF*, *SNRK*, and *PP2C*. *ARF* and *ERF* stand out as crucial TFs within the auxin as well as ethylene signaling pathways, respectively. By attaching to certain DNA sequences, they influence target gene transcription, thus impacting plant development and growth (Phukan et al., 2017; Cancé et al., 2022). Auxin has been shown to enhance FA synthesis in microalgae and increase monounsaturated FA levels (Jusoh et al., 2015; Dao et al., 2018). For research on wheat (Kovalevskaya, 2023) and Arabidopsis (Roudier et al., 2010), auxin is connected with the production of very long-chain FA. These very long-chain FAs play a function in plant reactions to stress from both abiotic and biotic sources (Batsale et al., 2021). On the other hand, *ERF* is also a key transcription factor when plants respond to stress (Müller and Munné-Bosch, 2015). Camellia fruit has a long development period, during which it is susceptible to abiotic stresses. Hence, these hormonal response factors significantly contribute to Camellia fruit’s growth and development. Moreover, recent findings have demonstrated ethylene’s capacity to boost the levels of linoleic acid and linolenic acid within *C. oleifera* fruit (Li et al., 2023). This also indicates that there is a complex relationship between plant hormones and plant lipids. Particularly under stress conditions, lipid remodeling emerges as a critical strategy for plant response to abiotic stressors (Chen et al., 2018; Liu et al., 2019). Studies have shown that spraying exogenous ABA on *C. oleifera* leaves under shortages of water significantly promotes the activity of genes such as *accA*, *accB*, *FAD6*, *FAD7*, and *FAD8* in the leaves (Yang et al., 2024). According to another study, the application of external ABA promoted the transcription of *FAD2* within oil palm fruit, which resulted in the accumulation of linoleic acid (Shi et al., 2021). This suggests the involvement of ABA in FA synthesis, with *SNRK* and *PP2C* as key elements in ABA signaling.

In the realm of agricultural practices, the strategic use of certain plant hormones has begun to show promising results in the growth and development of *C. oleifera* fruit (Li et al., 2023). This approach not only contributes to an increase in oil accumulation within the fruit but also aids in enhancing its overall quality. Such advancements highlight the potential of targeted hormone application in optimizing crop yields and improving the nutritional value of agricultural products. Given the intricate polyploidy and the wide array of *C. oleifera* varieties, there’s a pressing need for more in-depth research to uncover the molecular mechanisms behind how plant hormones influence oil accumulation in its fruit. Additionally, fine-tuning the application of these hormones and assessing their long-term impacts are essential measures to ensure the economic sustainability of agricultural practices. This dual approach will not only pave the way for maximizing oil yield and quality but also bolster the economic feasibility of cultivating *C. oleifera* on a larger scale.

## 5 Conclusion

This research undertook an in-depth investigation of the transcriptome and lipidome in *C. oleifera* seeds at the maturation stage, unveiling the FA accumulation patterns and their intricate association with gene expression regulation. The transcriptome analysis led to the identification of 26 344 genes that are expressed differently. Functional enrichment analysis indicated the involvement of phytohormone signaling transduction as well as starch and sucrose metabolism in oil accumulation. The notable high oil content in the kernel is largely due to the combined elevated expression of genes involved in the de novo fatty acid synthesis and triacylglycerol assembly pathways. Additionally, varietal expression variations in desaturases (*SAD*, *FAD*) result in differing unsaturated FA component levels among *C. oleifera* varieties. Particularly in high-oil varieties (HH2), the high expression of genes such as *FAD3*, *FAD7*, and *FAD8* during maturation increases the proportion of linolenic acid at the expense of the proportion of oleic acid. This reduction in oleic acid content is thought to encourage the synthesis of fatty acids through a potential feedback regulation mechanism, ultimately elevating the oil content in the kernel. In summary, the outcomes of this research offer a novel perspective on recognizing the molecular mechanism of oil biosynthesis in *C. oleifera* seeds, presenting valuable guidance for enhancing *C. oleifera* yields and improving tea oil quality.

## CRediT authorship contribution statement

**Dayu Yang:** Data Curation, Software, Formal analysis, Investigation, Visualization, Writing - Original Draft, Writing - Review & Editing**. Rui Wang:** Conceptualization, Methodology, Data Curation, Writing - Review & Editing, Supervision, Project administration**. Hanggui Lai:** Conceptualization, Methodology, Data Curation, Writing - Review & Editing, Supervision, Project administration**. Yongzhong Chen:** Conceptualization, Methodology, Writing - Review & Editing, Supervision. **Yimin He:** Investigation, Visualization, Data Curation. **Chengfen Xun:** Software, Data Curation, Validation. **Ying Zhang:** Data Curation, Validation. **Zhilong He:** Conceptualization, Methodology, Data Curation, Writing - Review & Editing, Supervision, Project administration, Funding acquisition

